# Epidemiological and ecological determinants of Zika virus transmission in an urban setting

**DOI:** 10.1101/101972

**Authors:** J Lourenço, M Maia de Lima, NR Faria, A Walker, MUG Kraemer, CJ Villabona-Arenas, B Lambert, E Marques de Cerqueira, OG Pybus, LCJ Alcantara, M Recker

**Affiliations:** Department of Zoology, University of Oxford, Oxford OX1 3PS, UK; Centre of PostGraduation in Collective Health, Department of Health, Universidade Estadual de Feira de Santana, Feira de Santana, Bahia, Brazil.; FIOCRUZ, Laboratory of Haematology, Genetics and Computational Biology, Salvador, Bahia, Brazil.; Institut de Recherche pour le Devéloppement (IRD), Universite de Montpellier, Montpellier, France.; Centre for Mathematics and the Environment, University of Exeter, Penryn Campus, Penryn TR10 9FE, UK

## Abstract

Zika has emerged as a global public health concern. Although its rapid geographic expansion can be attributed to the success of its *Aedes* mosquito vectors, local epidemiological drivers are still poorly understood. The city of Feira de Santana played a pivotal role in the early phases of the Chikungunya and Zika epidemics in Brazil. Here, using a climate-driven transmission model, we show that low Zika observation rates and a high vectorial capacity in this region were responsible for a high attack rate during the 2015 outbreak and the subsequent decline in cases in 2016, when the epidemic was peaking in the rest of the country. Our projections indicate that the balance between the loss of herd-immunity and the frequency of viral re-importation will dictate the transmission potential of Zika in this region in the near future. Sporadic outbreaks are expected but unlikely to be detected under current surveillance systems.

## Introduction

The first cases of Zika virus (ZIKV) in Brazil were concurrently reported in March 2015 in Camaçari city in the state of Bahia [1] and in Natal, the state capital city of Rio Grande do Norte [2]. During that year, the epidemic in Camaçari quickly spread to other municipalities of the Bahia state, including the capital city of Salvador, which together accounted for over 90% of all notified Zika cases in Brazil in 2015 [3]. During this period, many local Bahia health services were overwhelmed by an ongoing Chikungunya virus (CHIKV, East Central South African genotype) epidemic, that was introduced in 2014 in the city of Feira de Santana (FSA) [4, 5]. The role of FSA in the establishment and subsequent spread of CHIKV highlights the importance of its socio-demographic and climatic setting, which may well be representative of many other urban centres in Brazil and around the world, for the transmission dynamics of arboviral diseases.

On the 1^*st*^ February 2015 the first ZIKV cases were reported in FSA, followed by a large epidemic that continued into 2016. The rise in ZIKV incidence in FSA coincided temporally with an increase in cases of Guillain-Barré syndrome (GBS) and microcephaly [3], with an unprecedented total of 21 confirmed cases of microcephaly in FSA between January 2015 and December 2016. Although the causal link between ZIKV and severe manifestations is still under debate [6, 7, 8], the possible association has led to the declaration of the South American epidemic as an international public health emergency by the World Health Organization (WHO); the response to which has been limited to vector control initiatives and advice to delay pregnancy in the affected countries [9, 10]. With few cohort studies published and the lack of an established experimental model for ZIKV infection [11, 12], modelling efforts have taken a central role for advancing our understanding of the virus’s epidemiology [13, 14, 15, 16, 17, 18, 19]. In particular, quantifications of public health relevant parameters, such as the basic reproduction number (*R*_0_), the duration of infection [14], attack and reporting rates [20], the risk of sexual transmission [21] and birth-associated microcephaly [22, 18] are predominantly due to studies using transmission models. Crucially, however, and despite their major importance on the epidemiological dynamics of other arboviral diseases, such as dengue [23, 24, 25, 26] and chikungunya[27, 28, 29], the effects of local climate variables, such as temperature and rainfall, have not yet been explored in relation to Zika transmission.

In this study, focusing on an urban centre of Brazil (Feira de Santana), we explicitly model the mosquito-vector lifecycle under seasonal, weather-driven variations. Using notified case data of both the number of suspected Zika infections and gestations with neurological complications, we demonstrate how the combination of high suitability for viral transmission and low detection rates resulted in an extremely high attack rate during the first epidemic wave in 2015. The rapid accumulation of herd-immunity significantly reduced the number of cases during the following year, when the disease was peaking elsewhere in the country. Projecting forward we find that the demographic loss of herd-immunity together with the frequency of reintroduction will dictate the risk of re-mergence and endemic establishment of Zika in this and similar geographic settings in the near future.

## Materials and Methods

### Demographic and socio-economic setting

Feira de Santana (FSA) is a major urban centre of Bahia, located within the state’s largest traffic junction, serving as way points to the South, the Southeast and central regions of the country. The city has a population of approximately 620.000 individuals (2015) and serves a greater geographical setting composed of 80 municipalities summing up to a population of 2.5 million. Although major improvements in water supply have been accomplished in recent decades, with about 90% of the population having direct access to piped water, supply is unstable and is common practice to resort to household storage. Together with an ideal (tropical) local climate, these are favourable breeding conditions for species of the *Aedes* genus of mosquitoes, which are the main transmission vector of ZIKV, CHIKV and DENV that are all co-circulating in the region [25, 30]. FSA’s population is generally young, with approximately 30% of individuals under the age of 20 and 60% under the age of 34. In the year of 2015, the female: male sex ratio in FSA was 0.53 and the number of births 10352, leading to a birth rate standard measure of 31 new-borns per 1000 females in the population.

### Climate data

Local climatic data (rainfall, humidity, temperature) for the period between January 2013 and 2016 was collected from the Brazilian open repository for education and research (BDMEP, Banco de Dados Meteorológicos para Ensino e Pesquisa) [31]. The climate in Feira de Santana is defined as semi-arid (warm but dry), with sporadic periods of rain concetrated within the months of April and July. Between 2013 and 2015, mean yearly temperature was 24.6 celsius (range 22.5-26.6), total precipitation was 856 mm (range 571-1141), and mean humidity levels 79.5% (range 70.1-88.9%).

### Zika virus notified case data

ZIKV surveillance in Brazil is conducted through the national notifiable diseases information system (Sistema de Informação de Agravos de Notificação, SINAN), which relies on passive case detection. Suspected cases are notified given the presence of pruritic maculopapular rash (flat, red area on the skin that is covered with small bumps) together with two or more symptoms among: low fever, or polyarthralgia (joint pain), or periarticular edema (joint swelling), or conjunctival hyperemia (eye blood vessel dilation) without secretion and pruritus (itching) [32, 33]. The main differences to case definition of DENV and CHIKV are the particular type of pruritic maculopapular rash and low fever (as applied during the Yap Island ZIKV epidemic [34]). The data presented in Figure 1 (for both Brazil and Feira de Santana) represents notified suspected cases and was collected from the SINAN repository. Here, we use the terms *epidemic wave* and outbreak interchangeably (but see [18]).

**Figure 1.**
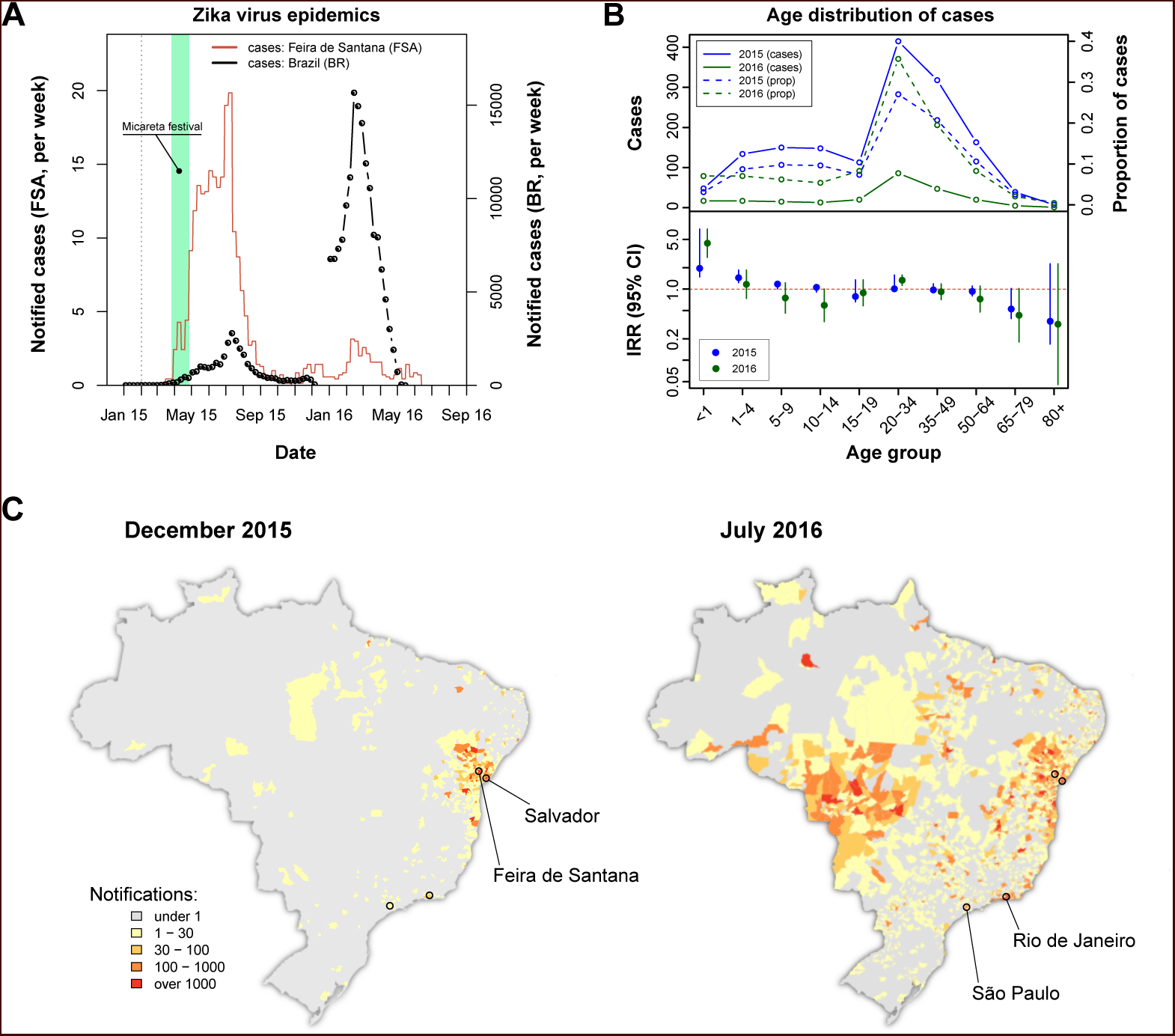
Zika virus epidemics in Feira de Santana and Brazil (2015-2016). *(A)* Comparison of weekly notified cases in Feira de Santana (FSA, full red line) and Brazil (BR, dotted black line). BR data for weeks 50-52 was missing. Green area highlights the time period for the Micareta festival and the dotted grey line the date of first notification. *(B)* Age distribution and incidence rate ratio (IRR) for the 2015 (green) and 2016 (blue) FSA epidemics. The top panel shows the number of cases per age (full lines) and the proportion of total cases per age class (shaded lines), which peak at the age range 20-50. The bottom panel shows the age-stratified incidence risk ratio (IRR, plus 95% CI), with the red dotted line indicating IRR=1. *(C)* Spatial distribution of cumulative notified cases in BR at the end of 2015 (left) and mid 2016 (right). Two largest urban centres in the Bahia state (Salvador, Feira de Santana) and at the country level (São Paulo, Rio de Janeiro) are highlighted.

### Microcephaly and severe neurological complications case data

A total of 51 suspected cases with microcephaly or other neurological complications were reported in FSA between January 2015 and December 2016. The first confirmed microcephaly case was reported on the 25th of October 2015 and virtually all consequent cases were notified before the summer of 2016. From these, after birth and follow up, 15 cases were cleared as not presenting abnormal development. Using guidelines for microcephaly diagnosis provided in March 2016 by the WHO (as in [35]), a total of 21 cases were confirmed by the end of 2016. Neurological complications not fitting microcephaly case definition were also found in 3 infants. A total of 3 foetal deaths were reported for mothers with confirmed ZIKV infection during gestation but for which no microcephaly assessment was available.

### Transmission model and fitting

To model the transmission dynamics of ZIKV infections and estimate relevant epidemiological parameters, we fitted an ento-epidemiological, climate-driven transmission model to ZIKV incidence and climate data of Feira de Santana between 2015 and 2016 within a Bayesian framework, similar to our previous work on a dengue outbreak in the Island of Madeira [23]. The general model structure is outlined below; full model details can be found as supporting material.

The model is based on ordinary differential equations (ODE) describing the dynamics of viral infections within the human and mosquito populations (eqn. S1-S5 and S6-S10, respectively). The human population is assumed to be fully susceptible before the introduction of ZIKV and is kept constant in size throughout the period of observation. After an infectious mosquito bite, individuals first enter an incubation phase, after which they become infectious to a mosquito for a limited period of time. Fully recovered individuals are assumed to retain life-long immunity. We assumed that sexual transmission did not significantly contribute to transmission dynamics and therefore ignored its effects [36].

For the dynamics of the vector populations we divided mosquitoes into two life-stages: aquatic and adult females. Adult mosquitoes were further divided into the epidemiologically relevant stages for arboviral transmission: susceptible, incubating and infectious. In contrast to human hosts, mosquitoes remain infectious for life. The ODE model comprised 8 climate-dependent entomological parameters (aquatic to adult transition rate, aquatic mortality rate, adult mortality rate, oviposition rate, incubation period, transmission probability to human, hatching success rate and biting rate), whose dependencies on temperature, rainfall and humidity were derived from other studies (see Table S1).

Four parameters (baseline mosquito biting rate, mosquito sex ratio, probability of transmission from human-to-vector and human lifespan) were fixed to their expected mean values, taken from the literature (see Table S2). To estimate the remaining parameters, alongside parameter distributions regarding the date of first infection, the human infectious and incubating periods, and the observation rate of notified ZIKV cases, we fitted the ODE model to weekly notified cases of ZIKV in FSA using a Bayesian Markov-chain Monte Carlo (MCMC) approach. The results are presented both in terms of mean dynamic behaviour of the ODE under the MCMC solutions and posterior distributions of key epidemiological parameters. A full description of the fitting approach and the estimated parameters can be found as supporting material.

## Results

On the 1^*st*^ February 2015 the first Zika virus (ZIKV) case was reported in Feira de Santana (FSA). Weekly cases remained very low for the following two months, adding up to just 10 notified cases by the end of March that year (Figure 1A). A rapid increase in the number of cases was observed in April, coinciding with Micareta, a local carnival-like festival that takes place across the urban centres of Bahia. The epidemic peaked in July 2015, which was followed by a sharp decline in notified cases over the next 1-2 months. This first epidemic wave was followed by a significantly smaller outbreak in 2016, peaking around March with only a handful of notified cases.

Overall, the epidemic behaviour in FSA was in sharp contrast with trends observed in notified cases across Brazil (BR), for which the second epidemic in 2016 was approximately 6 times larger than the one in 2015 (Figure 1A). Nonetheless, a clear temporal synchronization between country level and FSA case counts was observed, with the timing of first notifications in both datasets reiterating the Bahia state as a focus point in the emergence of ZIKV notified case data in Brazil [35, 3].

The age distribution of ZIKV notified cases in FSA suggested a higher proportion of cases between 20 and 50 years of age, but with no discernible differences between the two epidemics (Figure 1B, top panel). However, when corrected for the expected number of cases assuming an equal risk of infection per age class, we found the number of cases within this age group to be closer to most other groups (incidence rate ratio, IRR, close to 1, Figure 1B, bottom panel). The per capita case counts within the youngest age class (<1 years) appeared significantly more than expected, with an IRR significantly above 1 and also higher in 2016 (IRR=4.4, 95% CI [2.8, 7.0]) when compared to 2015 (IRR=1.95, 95% CI [1.5, 2.6]). There was also a consistent trend towards reduced IRR in the elderly (>65 years), although with significant uncertainty. Finally, a small increase in IRR was observed for the 20-34 year olds, which could potentially be a signature of sexual transmission in this age group [21, 30, 37, 38, 36]. With the lack of more detailed data it was not possible to ascertain whether these findings indicated notification bias, age-related risk of disease or simply age-dependent exposure risk, however.

The spatial distribution of total notified cases for BR highlighted the expected clustering of ZIKV cases within the Bahia state by the end of 2015 as well as the wider geographical range by July 2016 (Figure 1C). We speculated that the considerable difference in geographical range could explain the higher number of cases observed during the 2016 epidemic at the country level when compared to 2015. This, however, did not offer an explanation for why the second epidemic in FSA was nearly 7 times smaller than the first. To answer this question and to obtain robust parameter estimates of ZIKV epidemiological relevance we utilised a dynamic transmission model, which we fitted to notified case data and local climate variables of FSA within a Bayesian framework (see Materials and Methods).

## Climate-driven vectorial capacity

The reliance on *Aedes* mosquitoes for transmission implies a strong dependency of ZIKV transmission potential on temporal trends in the local climate. We therefore investigated daily rainfall, humidity and mean temperature data in FSA between 2013 and 2016 (Figure 2A). The data showed erratic fluctuations in rainfall with no clear seasonal trend and sporadic episodes of intense rain. Temperature, on the other hand, presented a much clearer seasonal signature with fixed amplitudes between 22 and 27 degree Celsius, peaking between December and May. Humidity showed an intermediate scenario and appeared correlated with periods of intense rainfall but negatively correlated with temperature.

**Figure 2.**
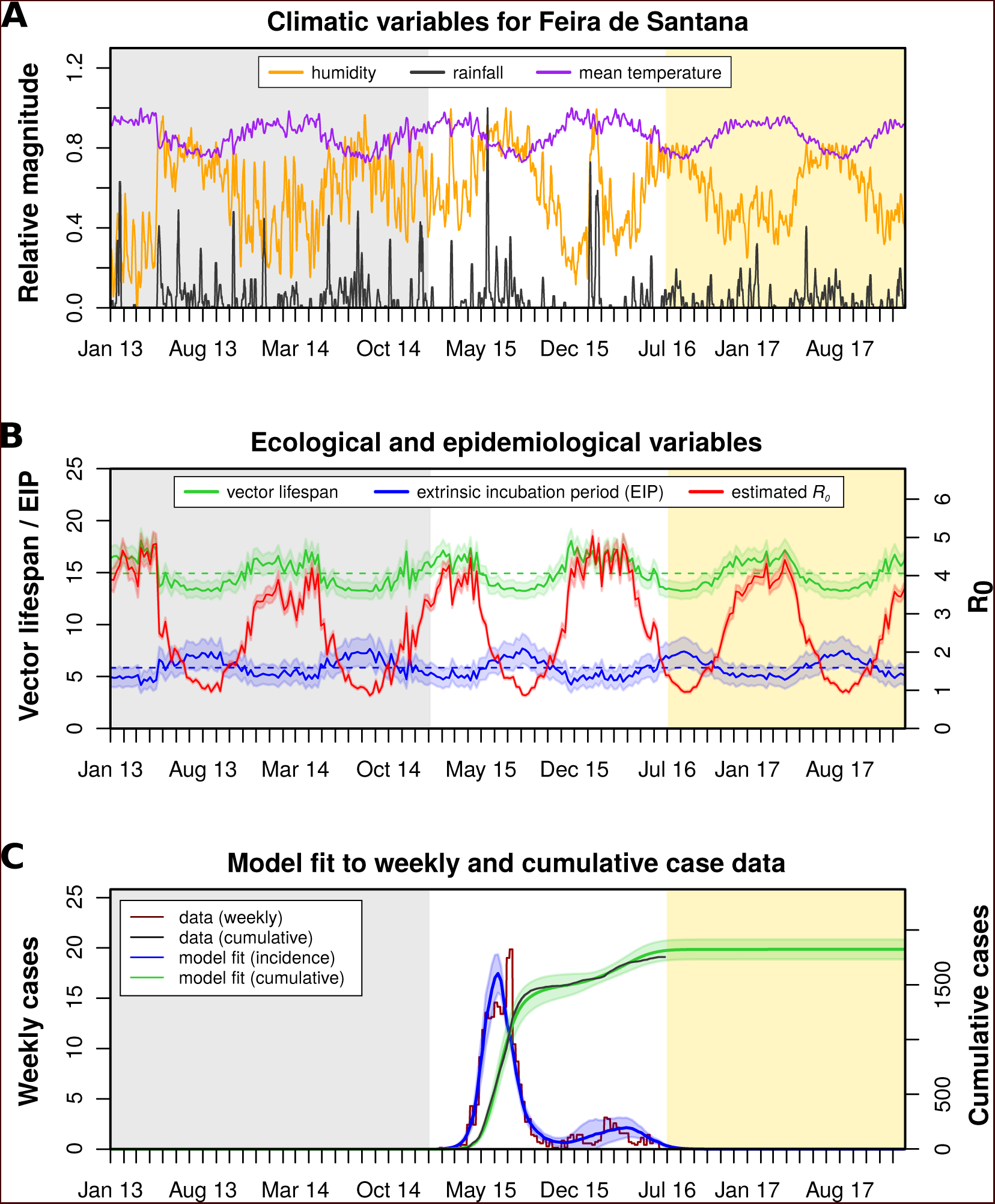
Eco-epidemiological factors and model fit to notified cases. *(A)* Daily climatic series for rainfall (black), humidity (orange) and mean temperature (purple) for Feira de Santana (FSA). *(B)* Estimated vector lifespan (green), extrinsic incubation period (EIP, blue) and basic reproduction number (*R*_0_, red). Median values are represented by horizontal dashed lines, with around 14.7days for the mosquito lifespan, 5.7 days for the EIP and 2.5 for *R*_0_ in the years preceding the epidemic (2.7 after 2015). *(C)* Resulting Bayesian MCMC fit to weekly (red line: data, blue line: model fit) and cumulative incidence (black line: data, green line: model fit). The grey areas highlight the period before the Zika outbreak, the white areas highlight the period for which Zika virus (ZIKV) notified case data was available, and the yellow shaded areas highlight the period for which mean climatic data was used (see Methods).

By fitting our weather-driven transmission model to these local climate and ZIKV case data we estimated the adult mosquito lifespan as well as the viral extrinsic incubation period (EIP) for the same period (see Materials and Methods). As shown in Figure 2B, both lifespan and incubation period showed seasonal oscillations with median values of around 14.7 and 5.7 days, respectively, which are well within the biological ranges found in the literature ([39, 40, 41, 42] and Table 1). Importantly, there was a strong negative temporal correlation between these two variables, with periods of longer EIP coinciding with shorter lifespans and vice-versa (Figure 2B).

**Table 1.**
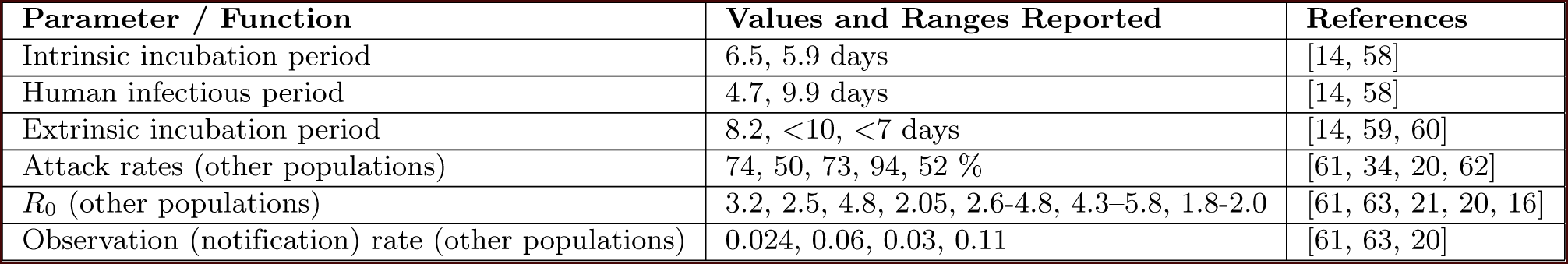
Literature-based reports on key ZIKV epidemiological and entomological parameters.

From these climate-driven parameters we next calculated Zika’s daily basic reproductive number, *R*_0_(*t*). As expected, the negative relationship between EIP and vector lifespan resulted in large temporal variations in vectorial capacity and thus seasonal oscillations in *R*_0_, with a median value of 2.5 (range [0.95, 4.35]), peaking in the local summer months between December and April. Following the qualitative climatic trends we found *R*_0_ to be highly correlated with mean temperature (*R*^2^ = 0.91) and humidity (*R*^2^ = 0.56) but not correlated with rainfall (*R*^2^<0.01). In agreement with other reports (Table 1), our estimated mean *R*_0_ for the two years preceding the ZIKV epidemic (2013-2014) was around 2.5. Notably, the transmission potential remained above 1 for the entire period, indicating a high suitability of viral transmission of ZIKV in FSA and hence other areas with similar climatic conditions.

## Model fit and parameter estimates

As shown in Figure 2C, our model closely reproduced both the dynamics and the sizes of the two epidemic waves in FSA (Figure 2C) between 2015 and 2016. We furthermore obtained a very close fit to the cumulative case data, which had reached a plateau by the end of the 2016 transmission season.

Four parameters of public health importance were estimated by our MCMC framework (Table S3): the date of introduction, the human infectious period, the human (intrinsic) incubation period, and the case observation rate. The posterior showed a strong support for an introduction in mid to late January, with an estimated median date of 14th of January 2015, i.e. three weeks before the first notified case (Figure 3A). Our estimated medians of the human incubation and infectious periods (Figures 3C, D) were 4.8 days (95% CI [2.6-9.8]).]) and 3.6 days (95% CI [2.2-5.0]) respectively, which were both within previously estimated ranges of ZIKV (Table 1).

**Figure 3.**
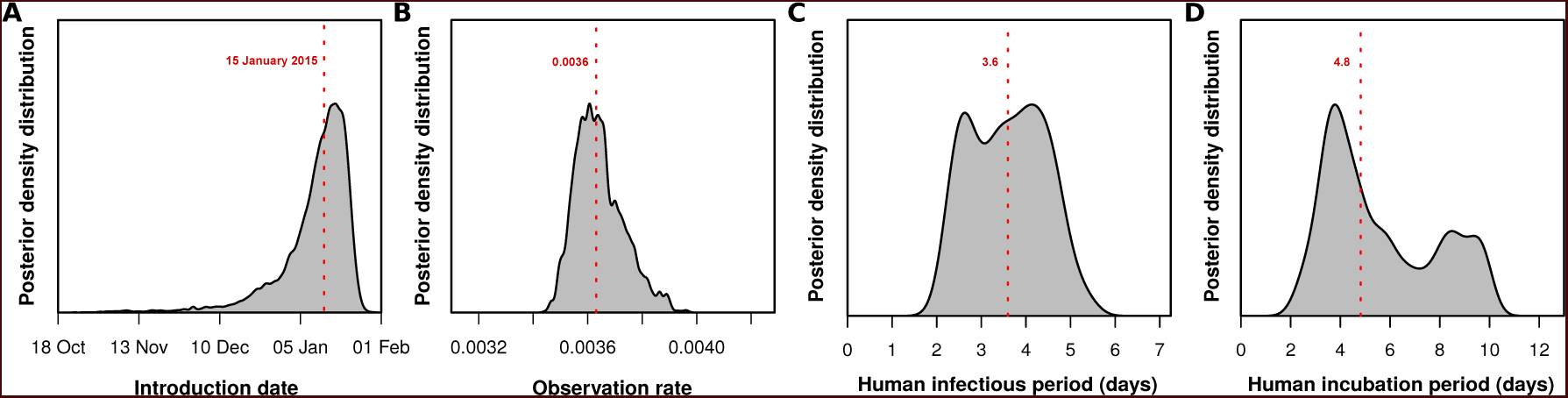
Estimated epidemiological and ecological parameters. Presented are the posterior distributions resulting from the MCMC fitting to FSA notified cases in 2015-2016, obtained from sampling 1 million MCMC steps. *(A)* Posterior for the introduction date with median 15^*th*^ January 2015 (95% CI [02/Jan/2015-27/Jan/2015]). *(B)* Posterior for the observation rate with median 0.0036 (95% CI [0.0035-0.0039]). *(C)* Posterior forthe human infectious period with median 3.6 days (95% CI [2.2-5.0]). *(D)* Posterior for the human (intrinsic) incubation period with median 4.8 days (95% CI [2.6-9.8]).

Of particular interest here was the very low observation rate (Figure 3B), with a mean of just 0.36% (95% CI [0.35-0.39]), which equates to less than 4 in 1000 infections having been notified during the epidemic in FSA. Although lower than other previous reported estimates, this would explain the relatively long period of apparently low viral circulation before the epidemic took off in April, 2015. That is, based on our estimates, there were just over 2,700 infections during the first 2 months, of which only 10 were notified. More importantly, when applying this rate to the total number of cases we found that by the end of the first epidemic wave around 69% (95% CI [66.3, 72.1]) of the population in FSA had been infected by the virus. This high attack rate is not unusual for Zika, however, and is in general agreement with observations elsewhere (Table 1). Furthermore, it also offered an explanation why the second wave in 2016 was so much smaller due to the substantial accumulation of herd-immunity during the first wave.

## Future transmission potential for Zika virus

As illustrated by the cumulative attack rate in Figure 4A, and similarly to estimates from other regions of the world (Table 1), nearly 70% of the population got infected by ZIKV infection by the end of 2015, which rose to over 80% (95% CI [77.7-85.9]) by the end of 2016. Notably, during the first wave most cases occurred off-season, here defined by our estimated daily reproductive number, *R*_0_(*t*), while the second wave appeared much more synchronized with the period of high transmission potential. Notably, this temporal phenomenon has also been observed for the chikungunya virus (CHKV) when it was first introduced into FSA in 2014 [5].

The amassed accumulation of herd-immunity during the first wave resulted in a marked difference between the estimated *R*_0_ and the effective reproductive ratio (*R*_*e*_) by the end of 2015 (Figure 4A), which in turn explained the huge reduction in Zika cases in FSA in 2016 when the virus was infecting a large number of individuals elsewhere (Figures 1A, C). By the beginning of 2016, *R*_*e*_ was estimated to be more than 3 times smaller than *R*_0_, which increased to 5 by the beginning of 2017. Projecting into the future using average climate data for this region and assuming normal demographic turnover in the population clearly showed that the effective reproductive number is expected to remain <1 for the next few years, suggesting a very weak potential for ZIKV endemicity in the near future.

**Figure 4.**
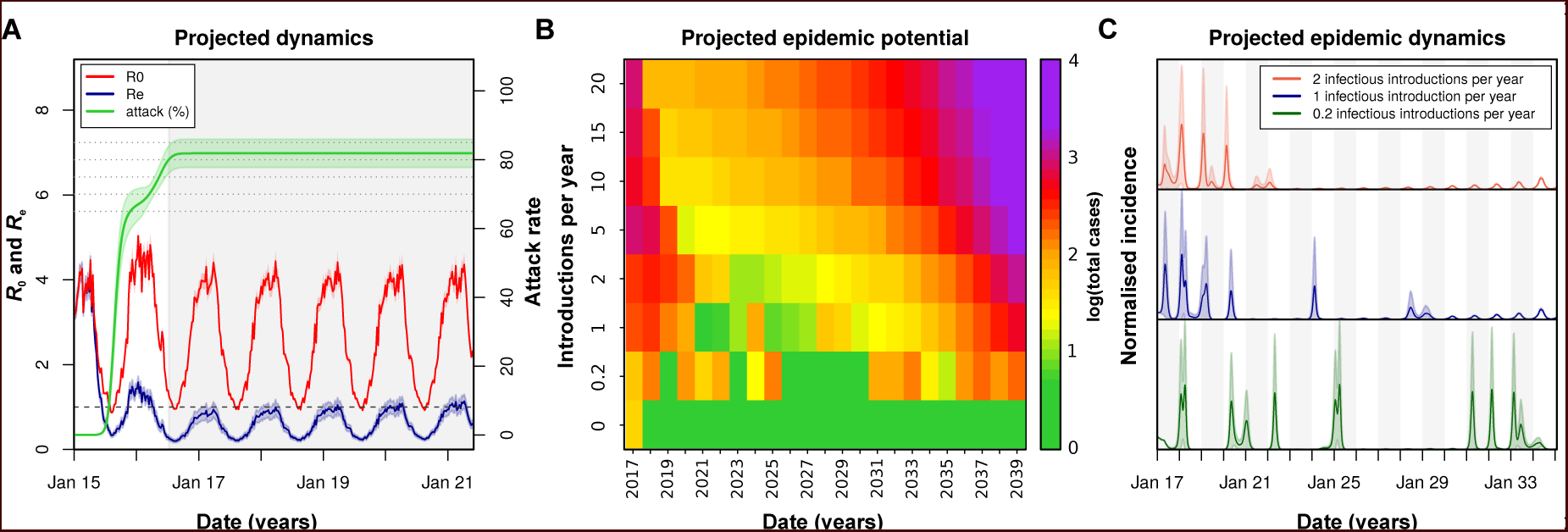
Projected Zika virus dynamics and transmission potential. *(A)* Fitted and projected epidemic attack rate (% population infected, green), basic reproduction number (*R*_0_, red) and effective reproduction number (*R*_*e*_, blue). Grey shaded area represents the period after the last available notified case. *(B)* Colourmap showing the projected total number of annual cases depending on rate of external introduction of infectious individuals. *(V)* Projected incidence dynamics when considering less than 1 (green), 1 (blue) and 2 (red) external introductions per year. Grey and white shaded areas delineate different years. The Y-axes are normalised to 1 in each subplot for visualisation purposes. In *(B, C)* results are based on 1000 stochastic simulations with parameters sampled from the posterior distributions (showing figure 3).

Without external introduction of infectious individuals (human or vector) our results suggested a high likelihood of an epidemic fade-out by 2017 (Figure 4A, B). We therefore projected ZIKV’s epidemic potential over the next two decades (until 2040) assuming different rates of viral introduction from other regions of Brazil or elsewhere (Figures 4B, C). In general, our results clearly showed that the potential for ZIKV to cause another outbreak or to establish itself endemically in FSA is strongly dependent on the frequency of re-introductions. In particular, the expected low transmission potential due to current herd-immunity levels together with infrequent introductions (≤ 1 a year) could result in long periods of no circulation followed by sporadic and unpredictable epidemics in the range of hundreds of infections (Figures 4B, C green line). In contrast, high rates of re-introductions could allow for semi-endemic behaviour but with periods of up to a decade presenting low incidence, during which susceptibility levels in the population would accumulate before ZIKV could cause larger epidemics. Notably, however, the levels of re-introduction required for endemic circulation may be unrealistically high (ex. 5-20), and we would argue instead that transmission of ZIKV in FSA will most likely be of sporadic and unpredictable nature in the years to come.

## Sensitivity to reporting and microcephaly risk

In effect, our observation rate entails the proportion of real cases that would have been notified if symptomatic and correctly diagnosed as Zika. Based on the previously reported Yap Island epidemic of 2007 [34], the percentage of symptomatic infections can be assumed to be close to 18%. Unfortunately, measures of the proportion of individuals seeking medical attention and being correctly diagnosed do not exist for FSA, although it is well known that correct diagnosis for DENV is highly imperfect in Brazil [43]. We therefore performed a sensitivity analysis by varying both the proportions of infected individuals seeking medical attention and the proportion of those being correctly diagnosed for Zika. Figure 5A shows that if any of these proportions is less than 10%, or with any combination of both less than 25%, our observation rate of 3.6 per 1000 infections can easily be explained.

**Figure 5.**
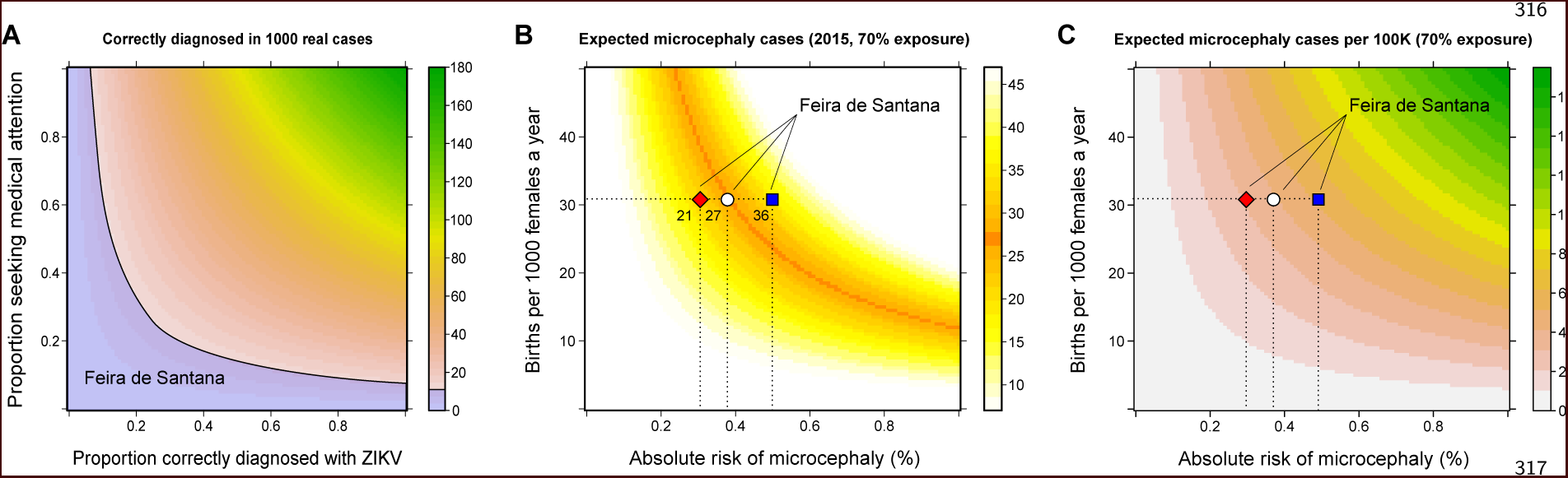
Sensitivity to reporting and microcephaly risk in Feira de Santana (FSA). *(A)* The observation rate (OR) can be expressed as the product of the proportion of cases that are symptomatic (0.18 [34]), with the proportion of symptomatic that seek medical attention, and the proportion of symptomatic that upon medical attention get correctly diagnosed with Zika. In the blue area the expected number of notified cases is 0-8 per 1000 real cases, a range with mean equal to FSA’s estimation (OR = 0.0036). *(B)* Expected number of cases of microcephaly (MC) and other neurological complications (NC) for theoretical ranges of birth rate (per 1,000 females) and risk of complications assuming 70% exposure of all pregnancies as estimated by our model for 2015 in FSA. *(C)* Expected number of MC and NC per 100,000 individuals under the same conditions as in *B*. The symbols in *B* and *C* represent the total confirmed MC cases (21, red diamond), the 21 MC plus 3 with NC and 3 fetal deaths (27, white circle), and 27 plus 9 currently under observation (36, blue square); the dashed horizontal line marks the number of births for FSA in 2015 (see Materials and Methods).

Finally we investigated the sensitivity of our results with regards to the expected number of newborns presenting either microcephaly or other neurological complications. Following the observation that virtually all reported cases were issued before the summer of 2016, we assumed that, if indeed associated with ZIKV infection during pregnancy, these would have been a consequence of the first epidemic wave in 2015. We therefore used the estimated attack rate of approximately 70% from 2015 (Figure 4A) and varied the local birth rate and the theoretical risk of ZIKV associated neurological complications to obtain an expected number of cases. In agreement with other reports [7, 44, 45, 46], our model predicted a relatively low risk for complications given ZIKV infection during pregnancy (Figures 5B, C and S2). In particular, using a conservative total of 21 confirmed microcephaly cases in FSA between 2015 and 2016, i.e. rejecting suspected or other complications, we estimate an average risk of approximately 0.3% of pregnancies experiencing ZIKV infection to develop microcephaly-associated syndromes. Including the 3 foetal deaths where ZIKV infections were confirmed during pregnancy plus 3 other cases with confirmed neurological complications after birth (total of 27 cases) increased the risk close to 0.4%, which rose further to 0.5 % when also including 9 individuals that are currently under surveillance.

## Discussion

Using an ento-epidemiological mathematical model of Zika virus (ZIKV) transmission, driven by temporal climate data and fitted to notified case data, we analysed the 2015-2016 outbreak in the city of Feira de Santana (FSA), in the Bahia state of Brazil. Using this framework we explored the epidemiological and ecological context of this outbreak and determined the conditions that led to the spread of the virus as well as its future endemic and epidemic potential. As FSA presents high suitability for Zika’s mosquito-vectors and with its particular geographical setting acting as a state commerce and transport hub, our results should have major implications for other urban centres in Brazil and elsewhere.

The epidemic pattern of ZIKV in FSA was in clear contrast to country-level observations, where case numbers were considerably higher during the second wave in 2016. In order to resolve whether this was due to a lower transmission potential of ZIKV in 2016 in FSA we calculated the daily reproductive number, *R*_0_(*t*), between 2013 and 2016. We found no notable decrease in 2016 in comparison to other years, but did notably find 2015 to present the highest potential, in line with the hypothesis that a particularly warm and wet year driven by El Niño may have temporarily boosted transmission potential of arboviruses [47]. By fitting our model to weekly case data we also estimated the observation rate, i.e. the fraction of cases that were notified as Zika out of the estimated total number of infections. It has previously been reported that the vast majority of Zika infections go unnoticed (Table 1), which is in agreement with our estimates of an observation rate below 1%. Based on this, around 69% of the local population were predicted to have been infected by ZIKV during the first wave in 2015, which is in the same range as Zika outbreaks in French Polynesia (66%) [44] and Yap Island (73%) [34]. The accumulation of herd-immunity caused a substantial drop in the virus’s effective reproductive number (*R*_*e*_) and hence a significantly lower number of cases during the second wave in 2016. In the context of FSA, it is possible that the high similarity of case definition to DENV, the concurrent CHIKV epidemic, and the low awareness of ZIKV at that time could have resulted in a significant number of ZIKV infections being classified as either dengue or chikungunya. Furthermore, based on our analysis we would argue that the percentage of correctly diagnosed ZIKV infections must have been exceptionally low (<20%).

The age structure of ZIKV notified cases in FSA showed a higher than expected incidence risk ratio (IRR) for individuals under the age of 4 years and a lower than expected risk for individuals aged +50 years. This contrasts the observation during the Zika outbreak on Yap Island in 2007, where all age classes, except the elderly, presented similar attack rates [34]. We note here, however, that the Yap Island analysis was based on both a retrospective analysis of historical hospital records and prospective surveillance (serology, surveys). It is therefore possible that the signatures amongst the youngest and oldest individuals in FSA may reflect deficiencies and/or biases in local notified data. In particular, such signatures could emerge by both a rush of parents seeking medical services driven by a hyped media coverage during the ZIKV epidemic, and a very small proportion of the elderly seeking or having access to medical attention. We also found a small increase in IRR in the 20-34 years age group, particularly during 2016, which could be indicative of the small contribution of sexual transmission. Most of these observations are speculative, however, and more detailed data will be required to fully understand these age-related risk patterns. For instance, initiatives such as the ZIBRA Project [48, 35], which perform mobile and real-time sampling with portable genome sequencing, could prove to be essential for a retrospective and future analysis of the ZIKV epidemic in Brazil, especially in areas where high levels of herd-immunity will prevent large-scale circulation in the coming years.

The implicit consideration of climate variables as drivers of vector biology in our model framework allowed us to ascertain the relative roles of temperature, humidity and rainfall for the reproductive potential (*R*_0_ and *R*_*e*_) of ZIKV. Similar to other studies in temperate and tropical settings, we found that temperature, with its direct influence on mosquito lifespan, aquatic development and extrinsic incubation period, was the key driver of seasonal oscillations in the transmission potential [23, 49, 28, 24]. Rainfall, on the other hand, only played a marginal role and appeared to be a relevant player for arboviral transmission mainly in tropical regions subject to intense rain seasons, such as areas in South East Asia [50, 51]. The oscillations in *R*_0_ presented seasonal windows of maximum transmission potential during the summer months between December and April. This would imply local surveillance and mitigation strategies, such as vector control, should be focused during this period with special attention between December and January, for which the virus could maximize its transmission potential.

A phylogenetic analysis has proposed that the introduction of ZIKV into Bahia took place between March and September 2014, although without direct evidence for its circulation in FSA at that time [52]. Our estimated date of introduction showed support for a date in mid-January 2015, a few months after the proposed introduction into Bahia. However, the posterior presented a wide margin stretching back to October 2014, suggesting that some of the parameter space obtained by the Bayesian approach is compatible with ZIKV circulation between September and December 2014. A mid-January introduction of ZIKV also implies a three week lag before the first case was notified in FSA. Similar periods between the first notification and estimated introduction often represent the time taken to complete one or more full transmission cycles (human-mosquito-human) before a cluster of cases is generated of sufficient size for detection by passive surveillance systems [23]. The case data also shows a 2-months period after the first notification during which weekly case numbers remained extremely low. This long period was unexpected as persistent circulation of ZIKV could hardly be justified by the observed total of only 10 cases. Given our estimated observation rate, however, the true number of ZIKV infections during this time amounted to just over 2,700 actual cases. In April, the number of notifications increased rapidly, coinciding with the Micareta festival, which we argue may have played a role in igniting the exponential phase of the epidemic by facilitating human-vector mixing as well as a more rapid geographical expansion. Another interesting observation was that the 2015 ZIKV epidemic peaked approximately 3 months after the estimated peak in the virus’s transmission potential, whereas there was much higher synchrony during the second wave in 2016. The same behaviour has been described for the chikungunya outbreak in FSA in 2014-2015 and which has been attributed to the highly discordant spatial distributions between the first two epidemics. It is likely that similar [5] or other heterogeneities [53] were responsible for the observed differences in the Zika outbreaks, although unfortunately we do not have access to spatial data to explore this hypothesis further.

After calibrating our model to the 2015-2016 epidemic, we projected the transmission of ZIKV beyond 2017 using stochastic simulations and average climatic variables representative of typical yearly trends. Without the possibility of externally acquired infections, local extinction was highly likely by 2017 due to high levels of herd-immunity. According to our study, Zika’s reproductive potential (*R*_*e*_) is expected to remain below 1 for at least another 5 years, given the slow replenishment of susceptibles in the population through births. When explicitly modelling the importation of infectious cases, our projections for the coming decades corroborated the conclusions of previous modelling studies that suggest a weak endemic potential for ZIKV after the initial exhaustion of the susceptible pool [14, 20]. However, our simulations further showed that the future epidemic behaviour was strongly dependent on the frequency of re-introductions, where sporadic and unpredictable epidemics could still be in the order of hundreds of cases. Furthermore, given our estimated observation rate for the 2015-2016 epidemic, the current passive surveillance system is unlikely to detect the occurrence of such small epidemics. Efforts should therefore be placed to improve ZIKV detection and diagnosis in order to optimize the local reporting rates.

Human sexual and vertical transmission of ZIKV infections is an important public health concern, especially within the context of potential Zika-associated microcephaly and other neurological complications in pre-and neonatals. With a total of over 10,000 live births in 2015 in FSA, our crude estimate for the risk of complications per pregnancy given ZIKV infection was between 0.3 and 0.5%, depending on the number of suspected and/or confirmed cases, which translates to about 4-6 expected cases per 100,000 individuals. As discussed elsewhere [44], this risk is extremely low when compared to other known viral-associated complications, such as those caused by infections by cytomegalovirus (CMV) and the rubella virus (RV) [54, 55]. It is therefore crucial to reiterate that what makes the ZIKV a public health concern is not the per pregnancy risk of neurological complications, but rather the combination of low risk with very high attack rates. According to our study, about 70% of pregnant women in FSA would have been challenged with ZIKV, equating to just above 7100 gestations at risk. Other studies have reported that the risk for complications during the 1^*st*^ trimester of gestation is higher than the one estimated here. For example, in the French Polynesia (FP) outbreak [44], the risk associated with ZIKV infection during the 1^*st*^ trimester was 1%, while the overall, full pregnancy risk was 0.42%, similar to our FSA estimates. For the Yap Island epidemic, no microcephaly cases have been reported. With an estimated number of 24 births per 1,000 females (census 2000 as in [34]) and using an overall risk of approximately 0.4% per pregnancy, only between 0-3 cases per 100,000 individuals would have been expected. However, the island’s small population size (7391 individuals [34]) together with a general baseline of 0-2 microcephaly cases per 100,000 in many areas of the world (ex. Brazil, Europe) [46, 56, 57], would explain the absence of reported cases.

There are certain limitations to our approach, many of which could be revisited when more detailed data becomes available. For example, we assumed homogeneous mixing between human and mosquito hosts but it is possible that spatio-temporal heterogeneities may have played a role in FSA. Furthermore, we have curated and integrated functional responses of key entomological parameters to temperature, rainfall and humidity variation, which were originally reported for dengue viruses. Due to lack of reliable data we have also assumed a constant observation rate for the 2015-2016 outbreak, whereas it is likely that due to a greater awareness and surveillance efforts to ZIKV in 2016, the rate may have been higher during the second wave. Our fitting approach is also dependent on notified case data and it is possible that the reported cases are not representative of the initial expansion of the virus, which may have thwarted the obtained posterior for introduction date. Finally, our future projections for the endemic and epidemic potential of ZIKV are based on average climatic trends of past years and do not capture the occurrence of natural variation between years, in particular for years affected by major Southern American climate events, such as the El Niño(a) [47].

In this study we have addressed the local determinants of ZIKV epidemiology in the context of a major urban centre of Brazil. Our results imply that control and surveillance of ZIKV should be boosted and focused in periods of high temperature and during major social events. These factors could identify windows of opportunity for local interventions to mitigate ZIKV introduction and transmission and should be transferable to any area for which both temperature data and community event schedules are available. We further confirm that the high transmission potential of ZIKV in areas like FSA can lead to the exhaustion of the local susceptible pool, which in turn dictate the long-term epidemic and endemic behaviour of the virus. Depending on the rate of re-introduction, sporadic outbreaks can be expected, although these will be unlikely to result in a notable increase in the number of microcephaly cases due to their limited sizes and low risk per pregnancy. Nonetheless, these sporadic occurrences could still have important, local public health consequences, and we argue that much better diagnostics and thus reporting rates are required for local authorities to detect and respond to such events in the near future. Our integrated mathematical framework is capable of deriving key insights into the past and future determinants of ZIKV epidemiology, which should be applicable to the eco-epidemiological settings of other major urban centres of Brazil and elsewhere.

## Acknowledgments

JL and AW received funding from the European Research Council under the European Union’s Seventh Framework Programme (FP7/2007-2013) / ERC grant agreement no. 268904 – DIVERSITY. MR was supported by a Royal Society University Research Fellowship. The European Research Council under the European Union’s Seventh Framework Program (FP7/2007-2013)/ERC grant agreement no. 614725-PATHPHYLODYN funded OP. MUGK’s contribution was made possible by the generous support of the American people through the United States Agency for International Development Emerging Pandemic Threats Program-2 PREDICT-2 (Cooperative Agreement No. AID-OAA-A-14-00102). CJVA was supported by a fellowship from the Labex EpiGenMed, via the National Research Agency, Program for Future Investment and University of Montpellier [ANR-10-LABX-12-01]. BL received funding from the Engineering and Physical Sciences Research Council (EPSRC) in the UK. NRF was supported by a Sir Henry Dale Fellowship jointly funded by the Wellcome Trust and the Royal Society (grant number 204311/Z/16/Z). The funders had no role in study design, data collection and analysis, decision to publish, or preparation of the manuscript.

